# Genome-wide Association Study Reveals that *PvGUX1_1* is Associated with Pod Stringlessness in Snap Bean (*Phaseolus vulgaris* L.)

**DOI:** 10.1101/2022.02.23.481584

**Authors:** Zhiyuan Liu, Shuo Gao, Helong Zhang, Zhaosheng Xu, Wei Qian

**Author notes:** These authors contributed equally to this work (Zhiyuan Liu, Shuo Gao). **Correspondence:** Wei Qian.

## Abstract

The suture strings is a particularly important pod trait that determines the quality and texture of snap bean (*Phaseolus vulgaris* L.). The *St* locus on chromosome 2 has been described as a major locus associated with suture strings. However, the gene and genetic basis underlying this locus remain unknown. Here, we investigated the suture strings of 138 snap bean accessions across two years. A total of 3.66 million single-nucleotide polymorphisms (SNPs) were obtained by deep resequencing. Based on these SNPs, we identified a strong association signal on Chr02 and a promising candidate gene, *PvGUX1_1*. Further analysis revealed that the 2-bp deletion in exon of *PvGUX1_1* was significantly associated with stringlessness. Comparative mapping indicated that *PvGUX1_1* was a domesticated locus and diverged from *PvGUX1_2* during an early stage. Our study provides important insights into the genetic mechanism of suture string formation and useful information for snap bean improvement.

## Introduction

Snap bean (*Phaseolus vulgaris* L.) is a type of common bean that is harvested before the seeds mature and eaten as a vegetable. Immature snap bean pods are succulent and rich in protein, carbohydrates, vitamin C, vitamin K, and carotenoids (Myers and Kmiecik, 2017). Therefore, the whole pods of snap bean are used for cooking, or preserved for freezing and canning (Hagerty et al.,, 2016). The snap bean is mainly consumed in North America, Europe, the Middle East, Africa, and Asia. In recent year, China has become the first producer of snap beans in the world (Zhang et al., 2008).

Improving pod quality is a major objective for snap bean breeding. Some pod characteristics, including pod length, pod shape, spur length, and the absence or presence of suture strings, have made the snap bean more palatable than the dry bean (another type of common bean in which the mature seed is consumed). A snap bean with the straight, smooth pod, and lacks suture strings is preferred in the fresh market. The fiber string along the suture is usually discarded before being eaten. Thus, the absence of suture strings is more popular with consumers and facilitates the commercial processing of snap bean.

Reducing suture strings in snap bean is crucial, and the key to this effort lies in understanding the genetic basis of the formation and development of suture string. The inheritance analysis of suture strings revealed that stringlessness was governed by a dominant gene, *St*, in common bean (Prakken, 1934). Quantitative trait locus (QTL) analysis located the *St* gene on chromosome Pv02 in common bean (Koinange et al., 1996). Working with a recombinant inbred line derived from dry bean and snap bean, a strong QTL, PST2.2, was also found on Pv02, accounting for 32% of total genetic variation in a recombinant inbred line (Hagerty et al., 2016). As the reduction of suture strings and pod wall fibers commonly lead to pod indehiscence in common bean, the indehiscent gene *PvIND* (a homolog of the Arabidopsis INDEHISCENT gene, *IND)* mapped near the St locus was predicted to be the *St* gene. However, there was incomplete co-segregation between *PvIND* and the St locus and a lack of polymorphisms with dehiscent/indehiscent phenotypes, suggesting that *PvIND* was not the gene *St* (Gioia et al., 2012). Recently, a single QTL, qPD5.1-PV, determining pod indehiscence was identified on chromosome Pv05 (Rau et al., 2019). In the attempt to identify the candidate gene underlying the QTL, a BC4/F4 introgression line population was used to narrow down the QTL in a 22.5 kb region and identified *PvMYB26* was the best candidate gene based on mapping and gene expression pattern (Di Vittori et al., 2020). In addition, several genes or QTLs were also discovered to be associated with pod dehiscence, such as *PvPdh1* on chromosome Pv03, QTLs on Pv08, Pv05 and Pv09 (Parker et al., 2020).

The first common bean reference genome was published in 2014 (Schmutz et al., 2014), which made it possible to use different strategies to identify candidate genes and molecular markers for important agronomic traits. Genome-wide association study (GWAS) is an approach based on using the numbers of single-nucleotide polymorphisms (SNPs) to test the association of desired traits. Due to the reduced cost of resequencing, and the repeatability of SNPs in the genome, GWAS has been performed using various landraces and breeding lines in common bean. These studies have focused on grain yield (Kamfwa et al., 2015; Moghaddam et al., 2016; Wu et al., 2020), flowering time (Raggi et al., 2019), resistance to disease (Wu et al., 2017), resistance to pod shattering (Parker et al., 2020), grain mineral content (Delfini et al., 2021), drought resistance (Wu et al., 2021), and abiotic stress (Soltani et al., 2018). However, few studies have focused on specific traits in snap bean (Myers et al., 2019). Pod stringlessness is particularly crucial in snap bean. Therefore, the objective of this study was to identify the candidate gene associated with this trait as a basis for further improving the quality of snap bean.

## Materials and Methods

### Plant material and resequencing

One hundred and thirty-eight snap bean accessions collected from the Institute of Vegetables and Flowers at the Chinese Academy of Agriculture Sciences (CAAS), including landraces, elite lines, and breeding lines, were grown between March and June in 2019 and 2020 (Supplementary Table 1). These seeds were grown in mixed nutrient soil at a greenhouse in Beijing(40° N, 116° E). The plants were watered using automatic drip irrigation every 2-3 days throughout entire growth period. The field away from plant was covered with a mulching plastic sheet to reduce the weed.

Young leaves at the unifoliate growth stage from each accession were collected, flash-frozen in liquid nitrogen and stroed in an ultra-low-temperatue freezer. Genomic DNAs were isolated for each genotype using Plant Genomic DNA kit (Tiangen, Beijing) following to the instructions. The integrity of gDNA was determined on 1% agarose gels. The concentration and quality of gDNA were measured using a NanoDrop2000 Spectrophotometer (Thermo Fisher Scientific). The DNA library were constructed accroding to the manufacturer’s instructions for the TruSeq nano DNA kit (Illumina). The libraries were genotyped using an Illumina HiSeq 2000 (125PE) sequencer at the facilities of Berry Genomics Co. Ltd, Beijing, China, as described by Wu et al. (2020).

### Measurement of pod sutures

At the green mature stage, 10 fresh pods from different plants of each accession were randomly chosen to measure the pod suture strings. The 10 pods from 10 individuals served as technical replications. The strings were evaluated as both a qualitative trait and a quantitative trait. As a qualitative trait, the pod strings were scaled 0–1 (0 = no suture strings, 1 = presence of suture strings). As a quantitative trait, the ratio (string length/pod length) of each pod was measured. The average ratio value of 10 pods and the scale rating of pods were both used for GWAS analysis.

### Expression analysis of *PvGUX1_1*

Three stages (T1 for pod elongation, T2 for pod development, and T3 for pod maturity) of R02 (non-suture pod) and R05 (suture pod) were sampled. The total RNA of the three stages of pods in suture pods and non-suture pods was extracted and converted to cDNA using a Reverse Transcription Kit (TransGen Biotech, Beijing, China) according to the manufacturer’s instructions. Quantitative real-time PCR was performed with SYBR Green (Vazyme Biotech, Nanjing, China), and the data collection was performed on QuantStudioTM 6 Flex system(ABI, Life, USA) according to the manufacturer’s instructions. The primers were synthesized by Sangon Biotech(Shanghai, China). The relative expression levels of *PvGUX1_1* were compared with that of β-actin, and the expression fold changes were calculated using the 2^-ΔΔCt^ method. Each qRT-PCR reaction was performed in triplicate. Sequences of the primers used for qRT-PCR in this study are shown in **Supplementary Table 2**.

### Variant calling and annotation

The raw paired-end reads were initially filtered by fastp (v0.20.0) software with the following parameters: -q 30. Next, the clean reads were aligned with the common bean reference genome V2.1 (Schmutz et al.,2014) using MEM algorithm of BWA (v0.7.17-r1188) (Li et al., 2009).

The tools SortSam and MarkDuplicates in PICARD (v1.127) were used to sort mapping results and mark the duplicate reads (https://broadinstitute.github.io/picard/). In addition, realignment around InDels was conducted by RealignerTargetCreator and IndelRealigner in GATK (v3.2) (McKenna et al., 2010). The variant was called by the UnifiedGenotyper module in GATK and SAMTOOLS (v1.6-3-g200708f) (Li et al., 2009). The two variant results were combined and further filtered to obtain a credible variant dataset using the GATK subcomponents SelectVariants and VariantFiltration. The credible variant dataset was employed to recalibrate and realign results using the BaseRecalibrator and PrintReads of GATK. The SNP and InDel were again called against the recalibrated results. Finally, a vcf file including all samples and variants was generated and further filtered using vcftools software (0.1.15) with the following parameters: -max-missing 0.95 -maf 0.05 -min-alleles 2 -max-alleles 2 -recode -recode-INFO-all.

The functional annotation of variants was performed using the software ANNOVAR (Version:2017-07-17) (Wang et al., 2010).

### Population genetics analyses

To analyze the population structure, the reduced SNPs were employed based on the value of the correlation coefficient (r^2^), where SNPs with strong linkage disequilibrium (LD) (r^2^ > 0.2) within a 50-kb window were discarded using plink (v1.90b6) software with the following parameters: -indep-pairwise 50 10 0.2. To estimate the most optimal sub-population, a cross-validation procedure was conducted with ADMIXTURE (v1.3.0) runs from K= 2 to 16 (Alexander et al., 2011). A neighbor-joining tree of 138 snap bean accessions was constructed using Phylip 3.68 (Felsenstein et al., 1989) software based on a distance matrix. The bar plots of sub-populations and the phylogenetic tree were plotted using the itol website (https://itol.embl.de/).

### Linkage disequilibrium analysis

The correlation coefficient (r^2^) of pairwise SNPs within a 1000-kb window from all chromosomes were used to estimate LD decay, which was calculated and plotted using PopLDdecay software (Zhang et al., 2019). LDBlockShow software was used to calculate and display LD blocks in candidate regions (https://github.com/BGI-shenzhen/LDBlockShow).

### Genome-wide association analysis

The high-quality SNPs were used for GWAS analysis in the R package GAPIT (Tang et al., 2016). To reduce false positives and improve statistical power, the ‘Q+K’ approach was employed. The kinship matrix (K) was calculated with the default method in GAPIT. The significant threshold (–log_10_*P*) was Bonferroni-corrected as –log_10_*P*=7.86. The Manhattan plot was run by the CMplot package in R 3.6.1 (https://github.com/YinLiLin/CMplot).

### Analyses of collinearity and synteny

The genome sequence information of common bean (*Phaseolus vulgaris* V2.1) and cowpea (*Vigna angularis* V1.2) were downloaded from phytozome 13. The genomes of soybean (*Glycine max* 109) (www.plantgdb.org/XGDB/phplib/download.php?GDB=Gm) and pea (*Pisum sativum*) (https://urgi.versailles.inra.fr/Species/Pisum) were downloaded from public databases. The analysis of collinearity and synteny between the four legumes was implemented with MCScan (Python version) (https://github.com/tanghaibao/jcvi/wiki/MCscan-[Python-version]). The proteins with similarity with over 90% on PvGUX1_1 in common bean, soybean, cowpea, and pea were identified using BLASTP with an e value <1e^-5^. The neighbor-joining tree from the orthologue gene of *PvGUX1_1* was constructed using MEGA X (Kumar et al., 2018) with default parameters.

## Results

### Pod suture string phenotyping

The pod suture strings of 138 snap bean accessions were investigated based on rating and ratio (Figure 1). For rating, the presence of strings was defined as 1; the absence of strings was defined as 0. The rating were investigated in 2019 (**Figure 1B**) and 2020 (**Figure 1D**). A total of 60 accessions were stringless, whereas 78 accessions had suture strings in 2019 (ST2-2019) (**Figure 1B**). However, five accessions showed different ratings in 2020 (ST2-2020). For ratio, the average ratio values (string length/pod length) of 10 pods in each accession were measured in 2019 (ST1-19) and 2020 (ST1-20) (**Figures 1A,C**). The analysis of correlation for ratio showed that there was a significantly high correlation of 0.93 (*P* = 0.00015) between 2019 and 2020.

**Figure. 1.**
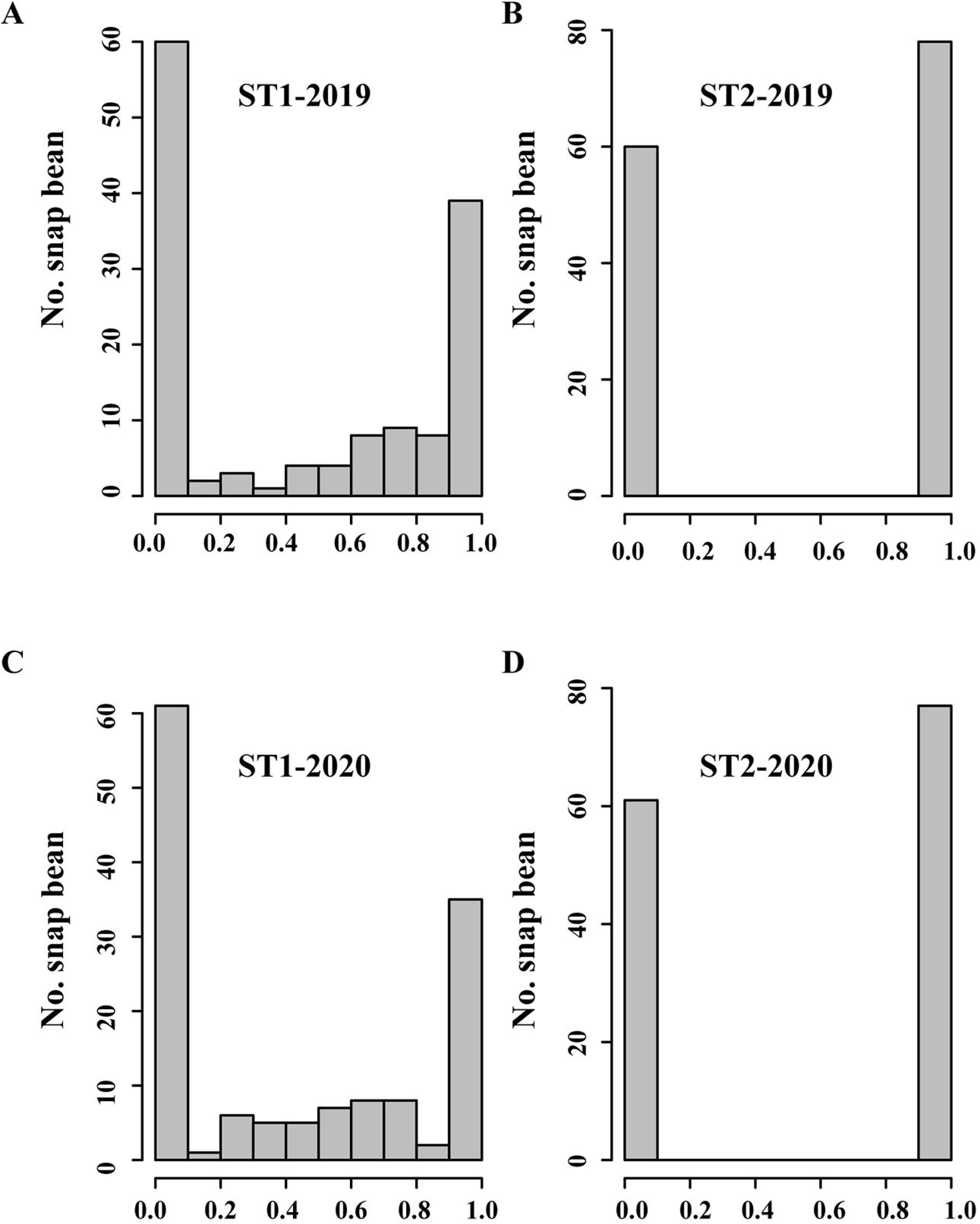
Histograms of pod suture strings in 138 snap bean accessions. (A) The ratio (string length/pod length) was measured in 2019. (B) The rating (1 for the presence of a string, and 0 for the absence of a string) was counted in 2019. (C) The ratio was measured in 2020. (D) The rating was counted in 2020.

### Resequencing of snap bean accessions

The whole-genome resequencing of 138 accessions produced a total of 3.08 billion raw paired-end reads and 0.92 Tb bases, which was approximately 11.4-fold sequence depth, ranging from 10.2- to 13.5-fold. After being filtered, 2.71 billion clean reads were retained. Mapping against the common bean genome V2.1 resulted in 5,130,030 SNPs and 1,524,528 InDels. Further filtering (bi-allelic, missing data < 0.05, minor allele frequency >0.05) identified 3,656,683 high-confidence SNPs and 626,853 InDels. Among these variants, 3,589,978 SNPs and 618,666 InDels were placed on chromosomes; 66,705 SNPs and 8187 InDels were on scaffolds. The distribution of these SNPs across the genome was uneven (**Figure 2**). Most SNPs were located in Chr02 (411,294), and the fewest SNPs were found in Chr06 (238,452). In addition, the frequency of SNPs in Chr02 (8.28 SNPs/kb) was the highest, while the frequency of SNPs in Chr08 (5.97 SNPs/kb) was the lowest **(Supplementary Table 3**).

**Figure. 2.**
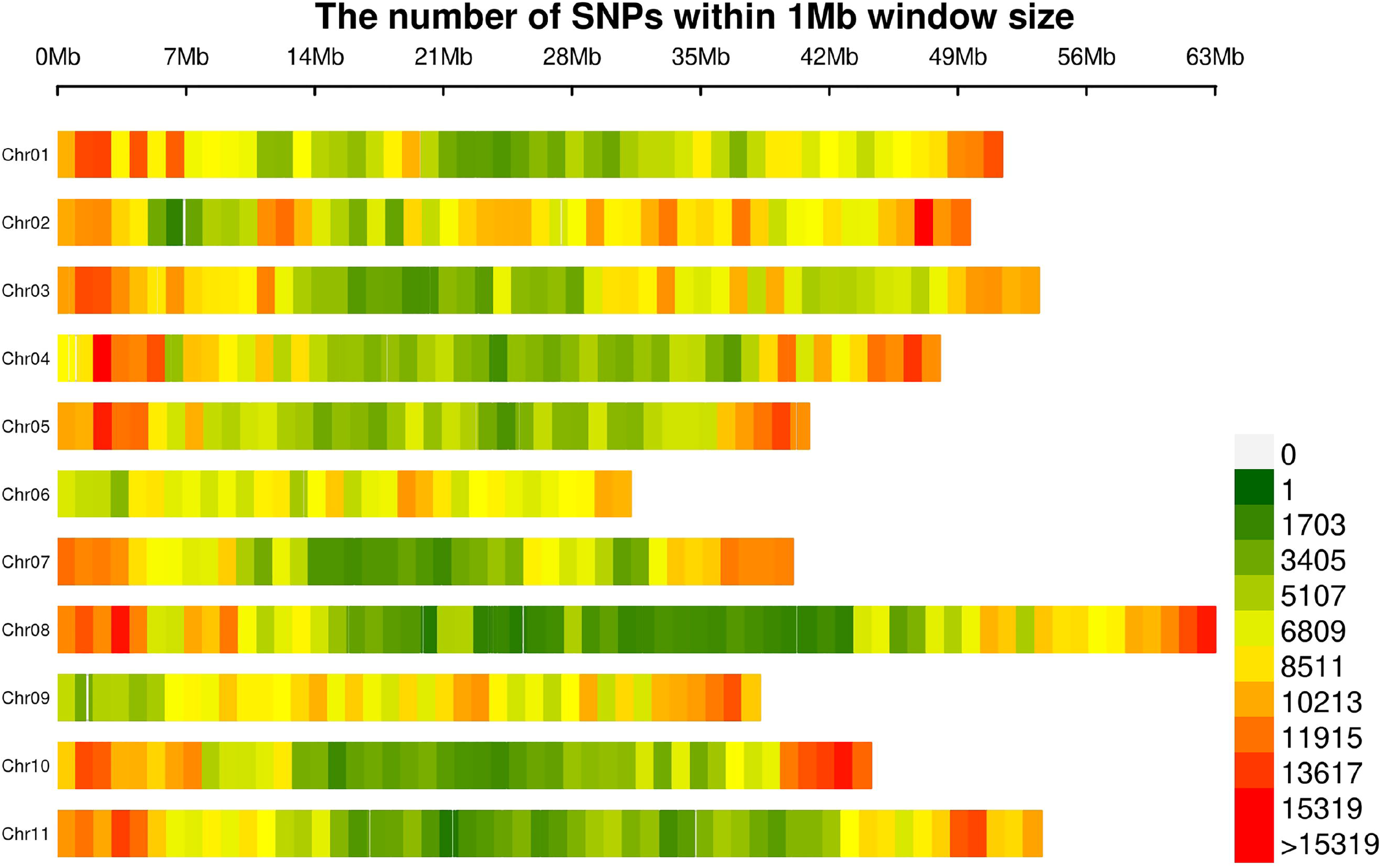
The number of SNPs within a 1-Mb window size across common bean chromosomes.

To investigate distribution regions of these variants across the genome, we carried out variant annotation and found that 146,882, 512,153, 279,102, and 244,805 SNPs and 4180, 53,742, 30,812, and 28,930 InDels were located in exons, introns, upstream, and downstream, respectively. Furthermore, of these SNPs in exons, 65,001 nonsynonymous, 718 stopgain, and 171 stoploss InDels were annotated, which resulted in amino acid changes, premature stop codons, or longer transcripts. Similarly, of these InDels in exons, 697 frameshift insertion, 1091 frameshift deletion, three stoploss, and 49 stopgain InDels were annotated, which also influenced protein sequences.

### Population structure and LD analysis

The analysis of population structure allows researchers to understand the genetic relationships and origins of species. After removing the SNPs with strong LD (r^2^ ≥ 0.2), 97,841 SNPs were generated and used to implement population structure analysis with Admixture. The use of K =2 divided the 138 genotypes into two genetic groups, which was in agreement with two domesticated genepools (Andean and Middle American) (**Figure 3**). Among the 138 genotypes, 40 genotypes had predominantly Andean ancestry, and 30 genotypes contained a level of hybridization, suggesting that a high degree of intercrossing between the genepools that has happened within snap beans.

**Figure. 3.**
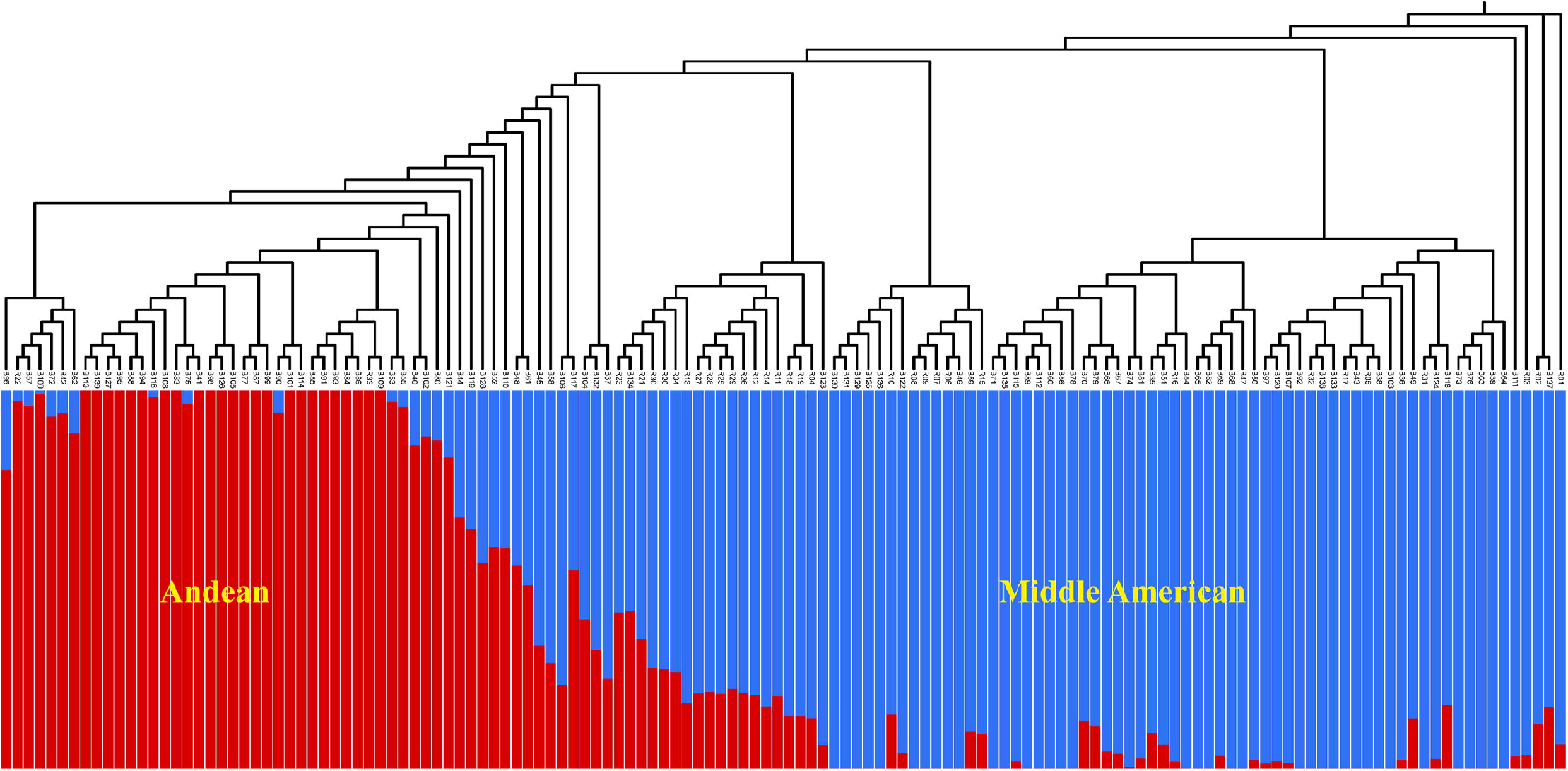
Neighbor-joining tree and population structure analysis using 97,841 single-nucleotide polymorphisms (SNPs). The genepools are colored with red for Andean and blue for Mesoamerican ancestry.

We further analyzed the LD decay across the genome **(Supplementary Figure 1**). The average LD decay of the whole genome was 631.4 kb (r^2^ dropped to half of its maximum value), which was faster than that of common bean (107 kb) (Wu et al., 2020), cultivated soybean (150 kb) (Zhou et al., 2015), and cultivated rice (123 kb for indica and 167 kb for japonica) (Huang et al., 2010). In addition, we found that the rate of LD decay in different chromosomes varied from 184 kb (Chr10) to 976 kb (Chr01) (**Supplementary Table 4**).

### Genome-wide association study for pod stringlessness

To find out genetic loci controlling pod stringlessness, we implemented GWAS for four traits (ST1-2019, ST2-2019, ST1-2020, and ST2-2020) using 3,656,683 SNPs (**Figure 4**). The Q2 and relatedness kinship matrix as covariates were taken into account to reduce false positives in GWAS analysis with a compressed mixed linear model. The –log10(*P*)=7.86 was set as a genome-wide significance threshold based on Bonferroni correction. Strong association signals were used to identify candidate regions and screen candidate genes.

**Figure. 4.**
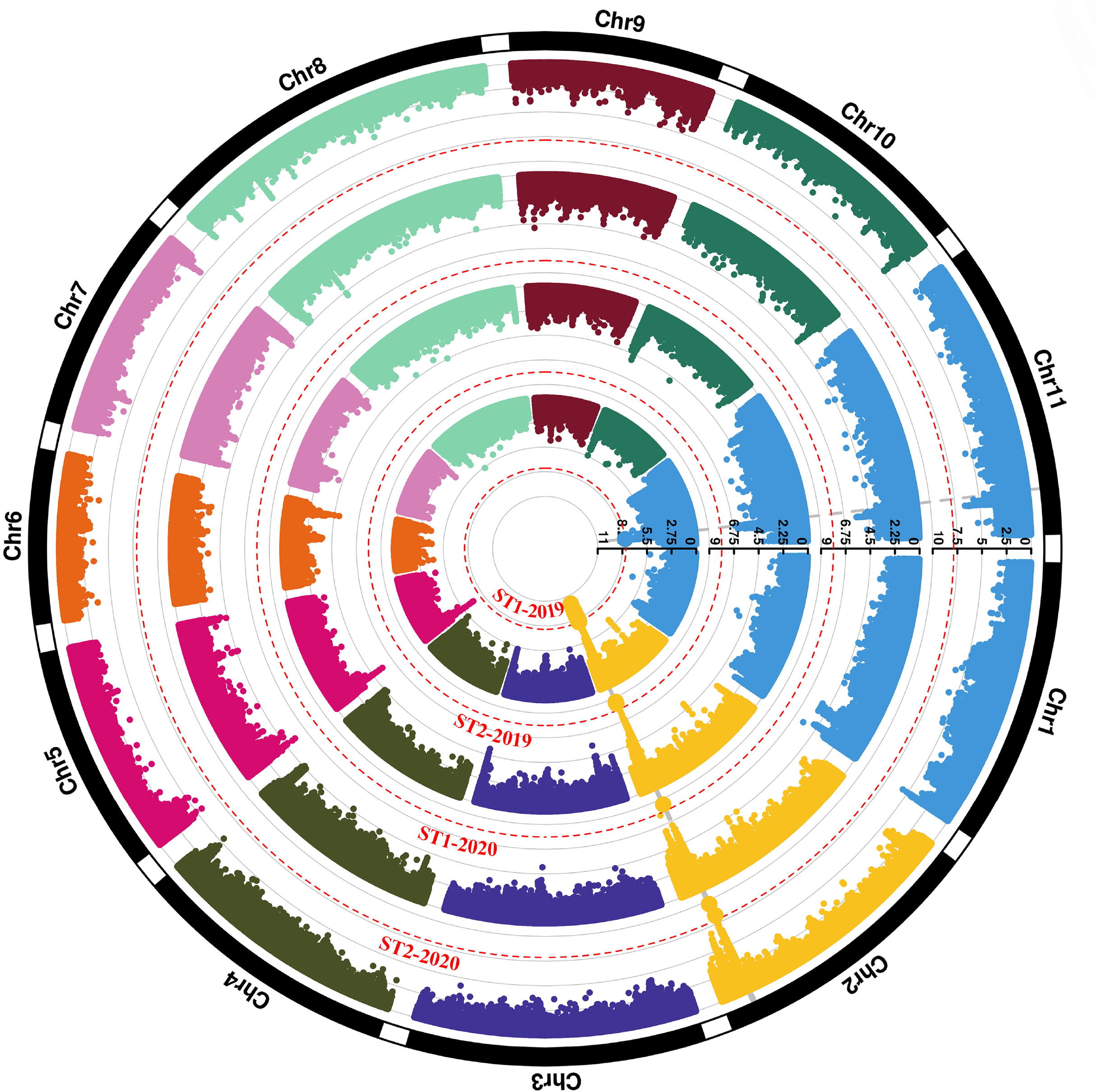
Circular Manhattan plots of a genome-wide association study (GWAS) for pod stringlessness. The inner circle to the outer circle represent ST1-2019, ST2-2019, ST1-2020, and ST2-2020, respectively. The red dashed lines of each circle indicate the threshold (7.86). Single-nucleotide polymorphisms (SNPs) over the threshold are highlighted.

A total of 205 loci were identified with –log10(*P*)>7.86 for ST1-2019. Of 205 SNPs, 204 were located at Chr02, and one was located at Chr11 (**Supplementary Table 5**). The peak signal was located at Chr02:44026689 (–log10(*P*)=10.08), accounting for 14.53% of phenotypic variation. The major locus Chr02:44248269 (–log10(*P*)=8.60) was significantly associated with ST2-2019. Furthermore, strong signals were both found at Chr02:44194640 for ST1-2020 (–log10(*P*)=8.49) and ST2-2020 (–log10(*P*)=9.62). Taken together, the peak SNPs for four traits were all located in adjacent physical regions in chromosome 2, which suggested the pod stringlessness was under the control of a major locus.

### Identification of candidate genes for pod stringlessness

To identify the candidate regions associated with the significant SNPs, we carried out haplotype analysis in the whole genome. We found that these peak SNPs for the four traits all resided on the same linkage disequilibrium (LD) block (Chr02:43998258–44264446) (**Figure 5**). These genes located in the block were likely responsible for the formation of stringlessness. **Table 1** shows these genes and their homologous genes in Arabidopsis. A total of 23 putative genes were annotated in this block based on the common bean reference genome V2.1. Eighteen out of 23 genes were functionally annotated, and 15 genes had homologous genes in Arabidopsis. Furthermore, 102 SNPs, including 43 nonsynonymous and 59 synonymous SNPs, and 6 InDels, including two frameshift deletions distributed in the coding areas of these genes, were also identified (**Table 2**).

**Figure. 5.**
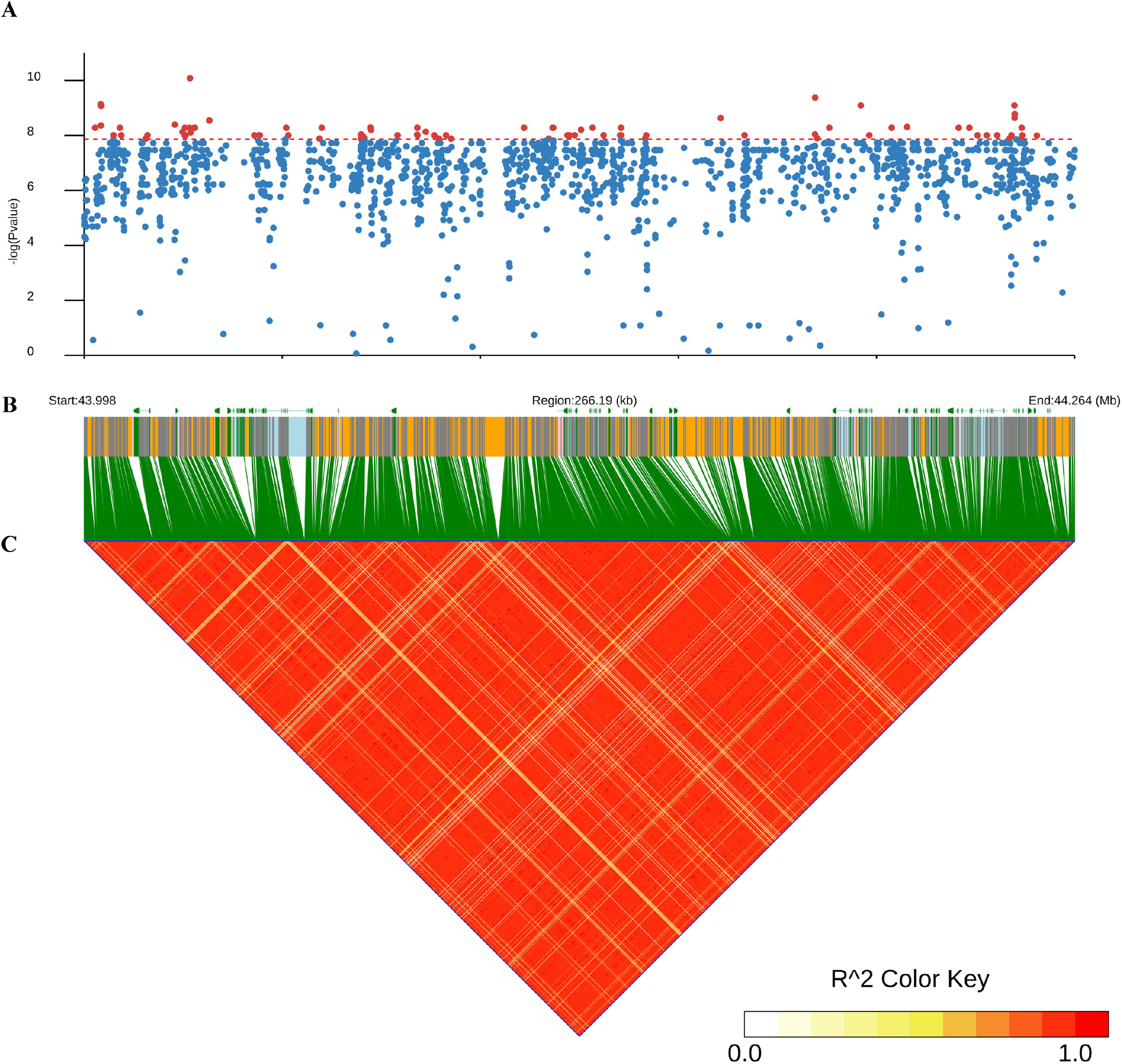
Manhattan plots and linkage disequilibrium (LD) heatmap over 266.29 kb around significant single-nucleotide polymorphisms (SNPs) on chromosome 2. (A) Manhattan plots of ST1-2019. The red dashed line represents the threshold (7.86). SNPs over the threshold are highlighted in red. (B) Annotated genes in the region. These CDS, introns, UTR, and intergenic regions are shown in green, light blue, pink, and orange, respectively. (C) The LD heatmap over the region. Colors are coded according to the r^2^ color key.

**Table 1.**
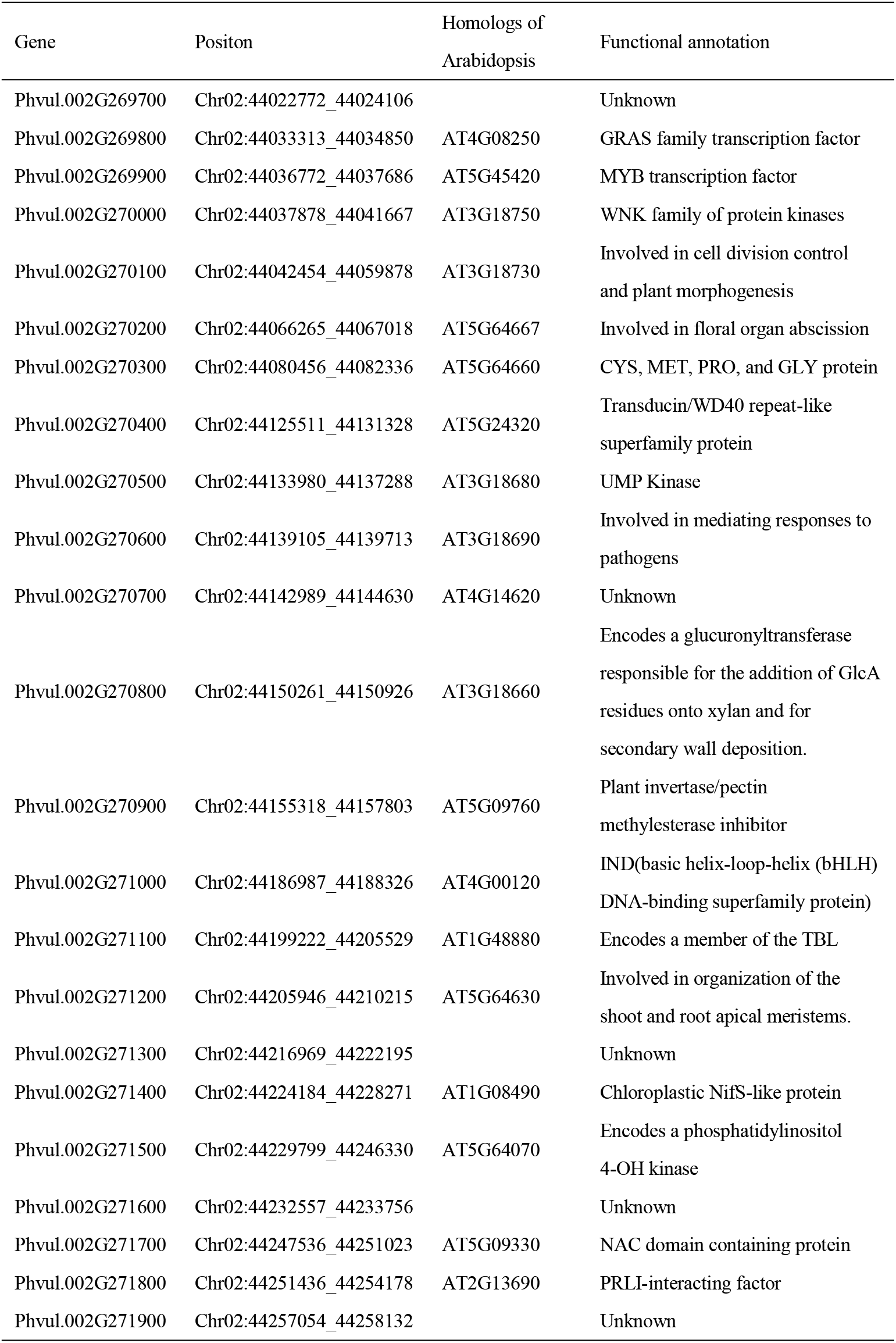
Putative genes in the 266.19 kb of candidate region

**Table 2.**
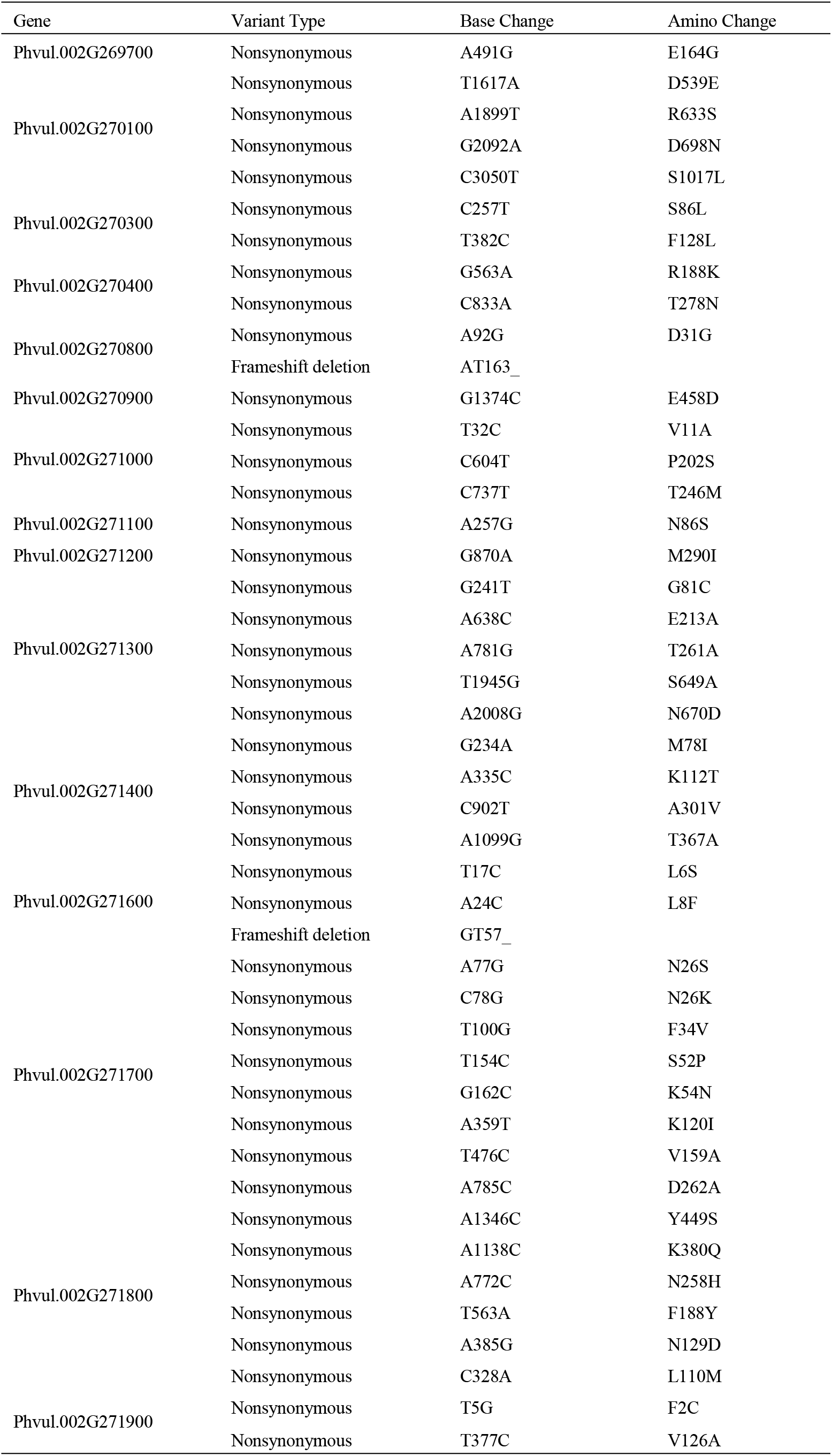
Functional annotation information of candidate genes

A 2-bp deletion in the exon region was identified in *Phvul.002G270800*, an ortholog to *AtGUX1*, which is responsible for secondary wall deposition in Arabidopsis. The 2-bp deletion introduced a premature stop codon that truncated the protein to 64 amino acids. To verify the deletion, we cloned the gene from two suture and non-suture accessions (**Supplementary Figure 2**). The result was similar to the finding in resequencing. Additionally, the deletion of 2 bp was significantly correlated with pod stringlessness (2.2×10^-16^) (**Figure 6**). We identified another a 2-bp deletion in gene *Phvul.002G271600;* however, the function of this gene was unclear.

**Figure. 6.**
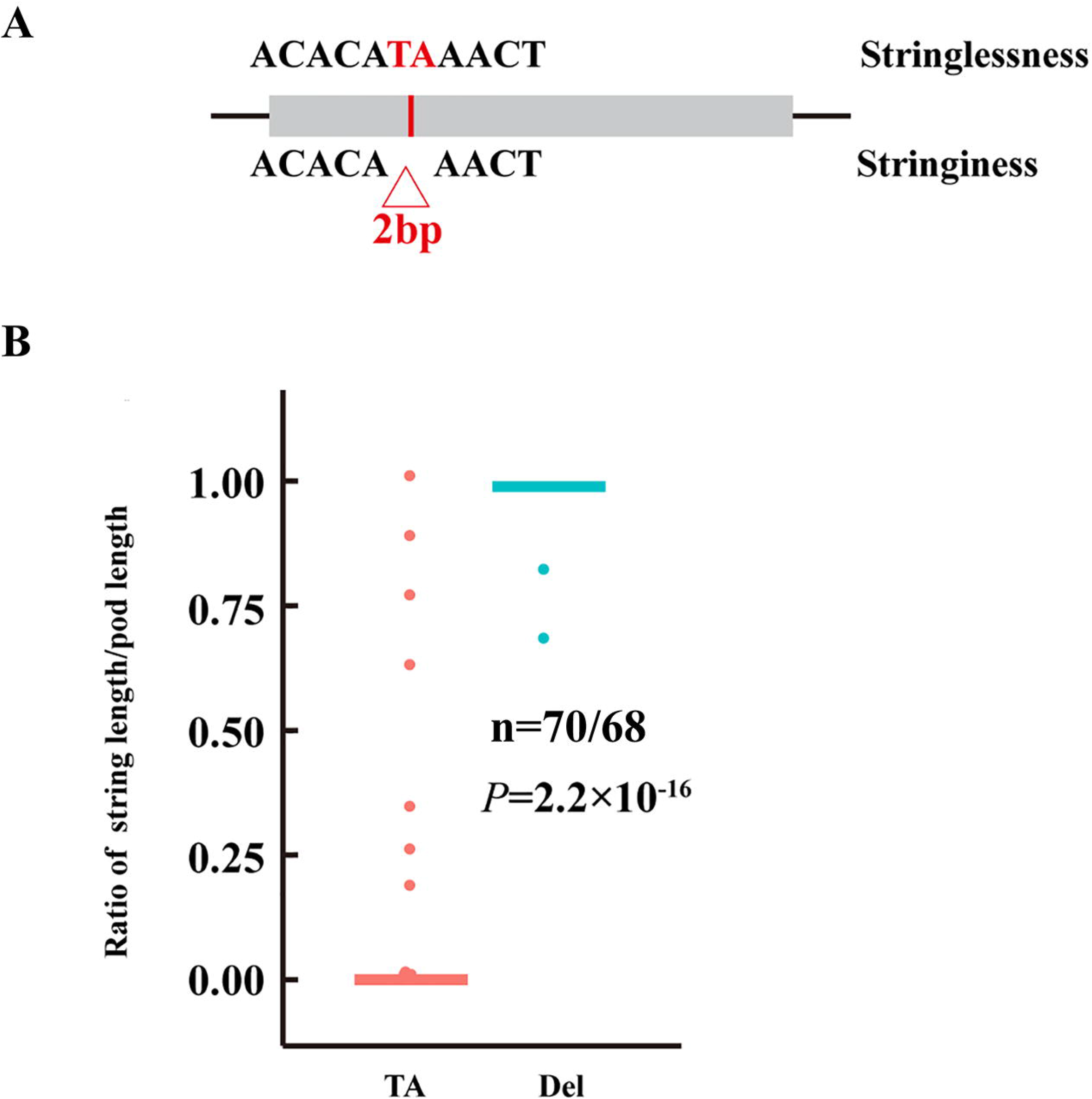
The identification of a 2-bp deletion in Phvul.002G270800. (a) Structure of Phvul.002G270800. The red base represents the 2-bp deletion in Phvul.002G270800. (b) Box plot of the ratio of string length/pod length for the 2-bp deletion in Phvul.002G270800.

The most abundant nonsynonymous SNPs were found in *Phvul.002G27l700. Phvul.002G271700*, encoding a NAC domain protein, carried eight nonsynonymous SNPs. Among these SNPs, K120I was significantly associated with pod stringlessness (1.39 ×10^-8^), whereas other SNPs exhibited weak association.

Three nonsynonymous SNPs, T32C, C604T and C737T, were identified in *PvIND (Phvul.002G271000)*. The SNPs T32C and C604 showed weak association (*P* = 6.42×10^-8^ and 6.79 ×10^-6^) with pod stringlessness, while C737T showed no association (*P* = 0.24).

### Syntenic analysis of the candidate region between the common bean and other legumes

To identify the function and relation of the candidate gene, we performed syntenic analysis within the candidate region of common bean with three legumes, including soybean (*Glycine max)*, cowpea (*Vigna angularis)*, and pea (*Pisum sativum)*. Common bean, cowpea, and soybean are members of the Phaseoleae tribe, whereas pea belongs to the Fabeae tribe (Dong et al., 2014). Amongst these legumes, the majority of cowpea are stringless, while common bean and pea have stringless and string types. In the Phaseoleae tribe, common bean and cowpea are the two most closely related crop species of the four legumes analyzed. They also exhibited a high degree of synteny (**Figure 7A**), in which 19 of 23 genes were orthologous. Although large-scale synteny with soybean was also observed, the homologous genes in soybean were divided into two regions (Glyma08g15530–Glyma08g15650 and Glyma08g16570–Glyma08g16640) on chromosome 8. In contrast, the pea chromosome exhibited a large rearrangement with common bean.

**Figure. 7.**
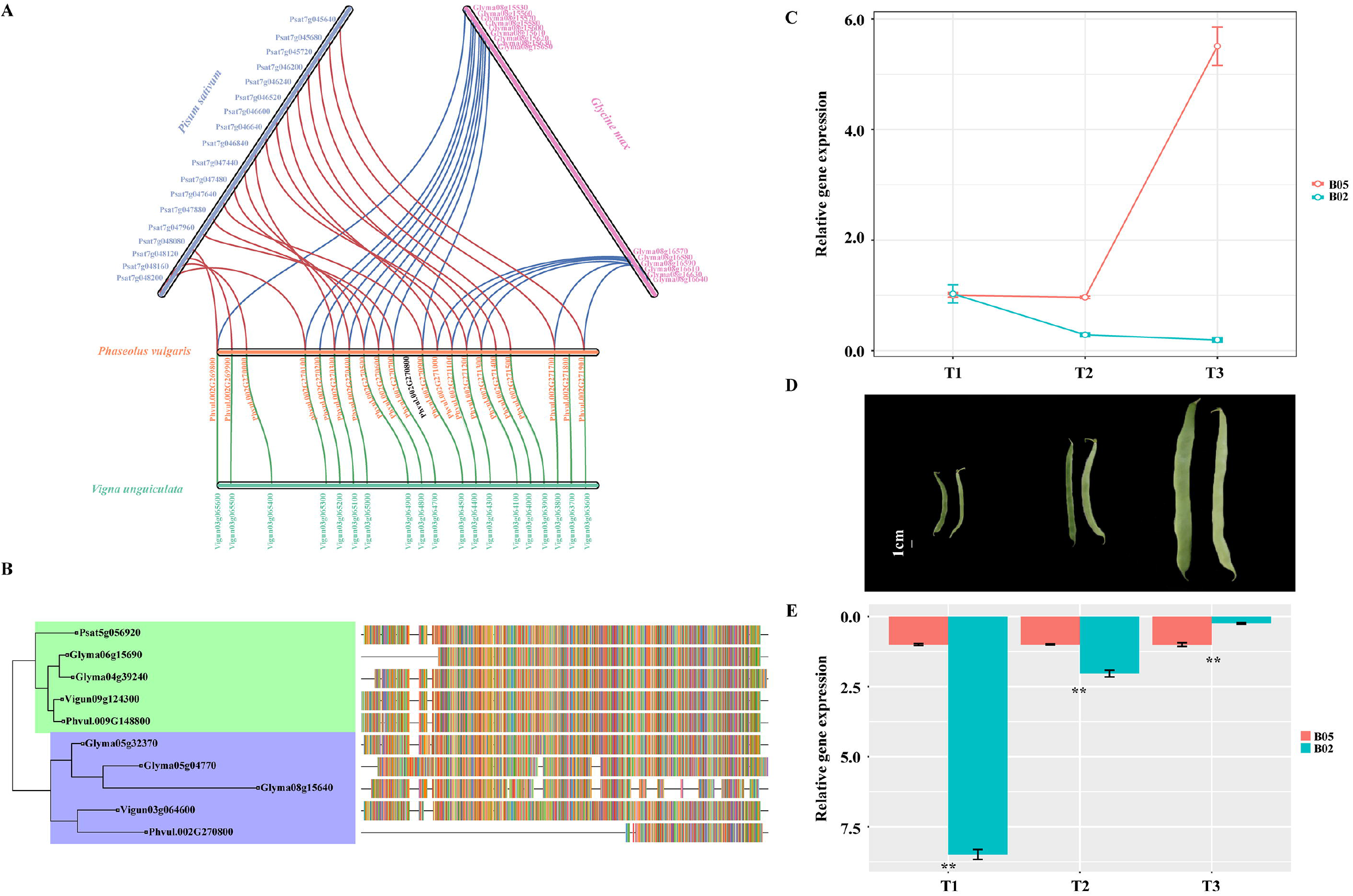
The analysis of phylogeny and expression of PvGUX1. A. Syntenic analysis of the candidate region between common bean and other legumes. B. Phylogenetic tree for PvGUX1. Colors located at the right side of each sequence represent their amino-acid composition. C. Gene expression of *PvGUX1_1* at different pod development stages for string and stringless snap bean. D. Morphology of stringless and string pod development stages T1–T3. E. The gene expression of PvGUX1_1 at three pod development stages. *P* values were calculated using student’s t test (* p < 0.05, ** p < 0.01).

Overall, these candidate genes and gene order in common bean were highly conserved and exhibited extensive synteny with cowpea. However, none of orthologs for Phvul.002G270800 in the syntenic block were identified (**Figure 7A**). To identify the orthologous gene of Phvul.002G270800 (PvGUX1_1), the protein of Phvul.002G270800 was used to conduct BLASTP search against cowpea, soybean, pea, and common bean. Specifically, we identified another common bean protein, Phvul.009G148800 (PvGUX1_2), which shared a strong sequence homolog to PvGUX1_1. PvGUX1_2 encoded 636 amino acids, whereas PvGUX1_1 encoded 221 amino acids. The best hit of PvGUX1_1 in cowpea, Vigun03g064600, encoded 629 amino acids, which exhibited large sequence difference with PvGUX1_1. To verify the relationship between PvGUX1_1, PvGUX1_2 and GUX1, we performed a phylogenetic analysis of PvGUX1_1, PvGUX1_2, and other orthologous genes. PvGUX1_1 and PvGUX1_2 were placed in two different sub-branches (**Figure 7B**). Although the corresponding genes, Glyma08g15640 and Vigun03g064600, in the synteny region were clustered with PvGUX1_1 in one sub-branch, there was large sequence variation between PvGUX1_1 and other orthologs. Collectively, these data demonstrated that PvGUX1_1 and PvGUX1_2 diverged at an early stage in legume evolution, which may have resulted in gene diversification.

### Gene expression patterns of PvGUX1_1

The formation of pod sutures is an important agronomic trait. To better reveal the genetic regulation of pod sutures, we conducted qRT-PCR analysis of PvGUX1_1 at the initiation of pod elongation (T1, no suture), pod development (T2, no suture), and pod maturation (T3, sutures were present in sutured pods) for sutured (R05)and non-sutured pods (R02) (**Figure 7D**). The qRT-PCR results indicated that the expression levels of PvGUX1_1 were significantly higher at the T1 and T2 stages in non-sutured pods compared with the sutured pods (**Figure 7E**). Furthermore, the expression level of PvGUX1_1 decreased following the development of pods in non-sutured pods (**Figure 7C**).

## Discussion

Understanding the genetic mechanism of suture string development will facilitate the study of domestication and plant breeding in snap bean. Here, we identified a strong signal on Chr02 that determined the formation of pod stringlessness based on large-scale resequencing. Within these putative genes in candidate regions, *PvGUX1_1* was the best candidate gene due to its function and polymorphisms, which was consistent with dominant inheritance.

### GWAS analysis for pod stringlessness

As common bean is a selfing species, effective recombination and heterozygosis in common bean are significantly reduced, which results in the generation of large LD and slow LD decay. Generally, LD decay is slower in selfing species than in outcrossing species because of the loss of recombination, which potentially leads to be homozygosity (Morrell et al., 2012). The nature of homozygosity makes common bean access to design GWAS. In particular, once a genotype is sequenced, the phenotype can be investigated in different environments. Moreover, the extensive genetic diversity is advantageous for GWAS analysis in common bean (Blair et al., 2009).

### Pod stringlessness was controlled by a major locus

The inheritance of pod stringlessness is complex due to genotype and environmental factors (Ma et al., 2016). Since the stringless trait was observed, various inheritance models for stringless trait in common bean have been proposed. Currence (1930) assumed that two genes (S for dominant, T epistatic to S) regulated the stringless trait. However, more studies revealed that the stringless trait was under the control of a single dominant locus (*St*), which was mapped on chromosome 2 (Koinange et al., 1996; Davis et al., 2006; Gioia et al., 2012). Moreover, there have also been reports that the trait did not fit the ratio of one or more loci, and thus was a quantitative trait (Hagerty et al., 2016). In order to verify the inheritance pattern, qualitative traits and quantitative traits were both used for GWAS analysis. Interestingly, we obtained similar results from the two models. The only strong signal from both models was identified on Chr02, which was in agreement with previous findings, and showed that the formation of suture string was controlled by a major locus.

The formation of suture string is controlled by a single locus, while the level (short versus long) of the string might be more complex. This characteristic was also observed in pod shattering. As suggested by Rau et al., (2019), at least two additional loci were likely relevant to the level and mode of pod shattering. In our study, in addition to Chr02, a SNP located at Chr11 was also associated with stringlessness (**Supplementary Figure 3**). The SNP was about 0.13 Mb from the NAC transcription factor gene *PvCUC2* (*Phvul.011G160400*, Chr11:45614432_45616861). In order to identify more locus, we conducted GWAS only on stringy snap bean for ST1-2019. Strong association signals were identified on Chromosome 7 (Supplementary Figure 4). These locus may be responsible for the level of suture string, along with the St gene. This finding supported the hypothesis by Drijfhout (1978) that a major factor influenced the string formation trait, while other genes led to incomplete strings.

### Candidate gene for stringlessness in snap bean

A total of 23 genes within the LD block surrounding the high association signal were identified. Among them, several genes are orthologous genes involved in cell-wall biosynthesis, pod shattering, and fiber formation. *Phvul.002G270800* is the orthologous gene of *AtGUX1* (*AT3G18660*). In Arabidopsis, AttGUX1 belongs to Glycosyltransferase Family 8, which participates in the synthesis of plant cell walls (Yin et al., 2010). *AtGUX1* is responsible for the decoration of xylan, an important component of the secondary cell wall (Lee et al., 2012). Silencing *AtGUX1* led to the decrease of glucuronoxylan content and microsomal xylan in the cell wall (Oikawa et al., 2010). In our study, a 2-bp deletion was found in the exon region of *Phvul.002G270800*, causing a premature stop. The 2-bp deletion was significantly associated with pod stringlessness. Therefore, we propose *Phvul.002G270800* as the best candidate gene for St locus.

In addition to *Phvul.002G270800*, another gene of interest was *Phvul.002G271000*, the orthologous gene to *AtIND. AtIND*, as a b-HLH transcription factor, plays a crucial role in pod shattering in Arabidopsis (Girin et al., 2010; Kay et al., 2013; Dong and Wang, 2015). However, due to the lack of mutation in *PvIND* (*Phvul.002G270800*), a previous study reported that it was not the *St* gene controlling suture strings (Gioia et al., 2012). Although the present study identified three nonsynonymous SNPs in the exon region of *PvIND*, these SNPs only showed a weak association with the suture strings. Therefore, *PvIND* may not be directly involved in suture development. Other interesting genes included NAC transcription factor *Phvul.002G271700* (*PvVNI1*) and MYB transcription factor *Phvul.002G269900 (PvMAMYB)*. Many studies have suggested that an NAC transcription factor is correlated with pod shattering and secondary cell wall development (Hussey et al., 2011; Yamaguchi et al., 2010; Reusche et al., 2012). In particular, the role of the NAC transcription factor SHA1-5 in regulating pod shattering in soybean has been elucidated in detail (Dong et al., 2014). Likewise, several MYB transcription factors, such as *MYB26* (Wilson et al., 2011), *MYB46* (Kim et al., 2013,2014), and *MYB63* (Zhou et al., 2009), are involved in lignin biosynthesis and secondary cell wall formation in many species (Nakano et al., 2015). Therefore, the functions of *PvVNI1* and *PvMAMYB* need to be further studied in future research to determine whether they are related to suture string development or pod shattering.

### Pod stringlessness in other legumes

The loss or presence of suture strings is not an important factor for many legumes in which the dry seeds are consumed. In legumes, reports on the stringless trait are currently only found in common bean and pea. In pea, pod stringlessness arose from spontaneous mutation (Wellensiek, 1971). The recessive gene (*sin-2*) is regarded as the key gene responsible for the stringless trait in pea (McGee and Baggett, 1992; Ma et al., 2016). In contrast, the stringless trait in common bean is governed by the dominant gene *St*. In the synteny block, the orthologs of *GUX1* were not detected in pea, indicating that the genetic mechanism of stringlessness between the two legumes may be different.

Although the same regulation gene may not be shared in common bean and pea, there are many parallels, including seed dormancy, growth habit, and earliness, between common bean and pea that have occurred in the process of crop domestication (Weeden, 2007). The identification of the *St* gene in common bean would accelerate the mining of *sin-2* and improve the understanding of the genetics of domestication under parallel selection in the future.

## Data Availability Statement

The original contributions presented in the study are included in the article/Supplementary Material, further inquiries can be directed to the corresponding author.

## Author Contributions

QW designed the study. LZ and GS conducted the experiments. LZ, ZH, and XZ analyzed the data. LZ wrote the manuscript. QW, XZ, ZH, and GS revised the manuscript.

## Supplementary Material

The Supplementary Material for this article can be found online.

