## Supplementary for "Genome-wide Association Study Reveals that *PvGUX1_1* is Associated with Pod Stringlessness in Snap Bean (*Phaseolus vulgaris* L.)": Supp figures1-4 .DOCX

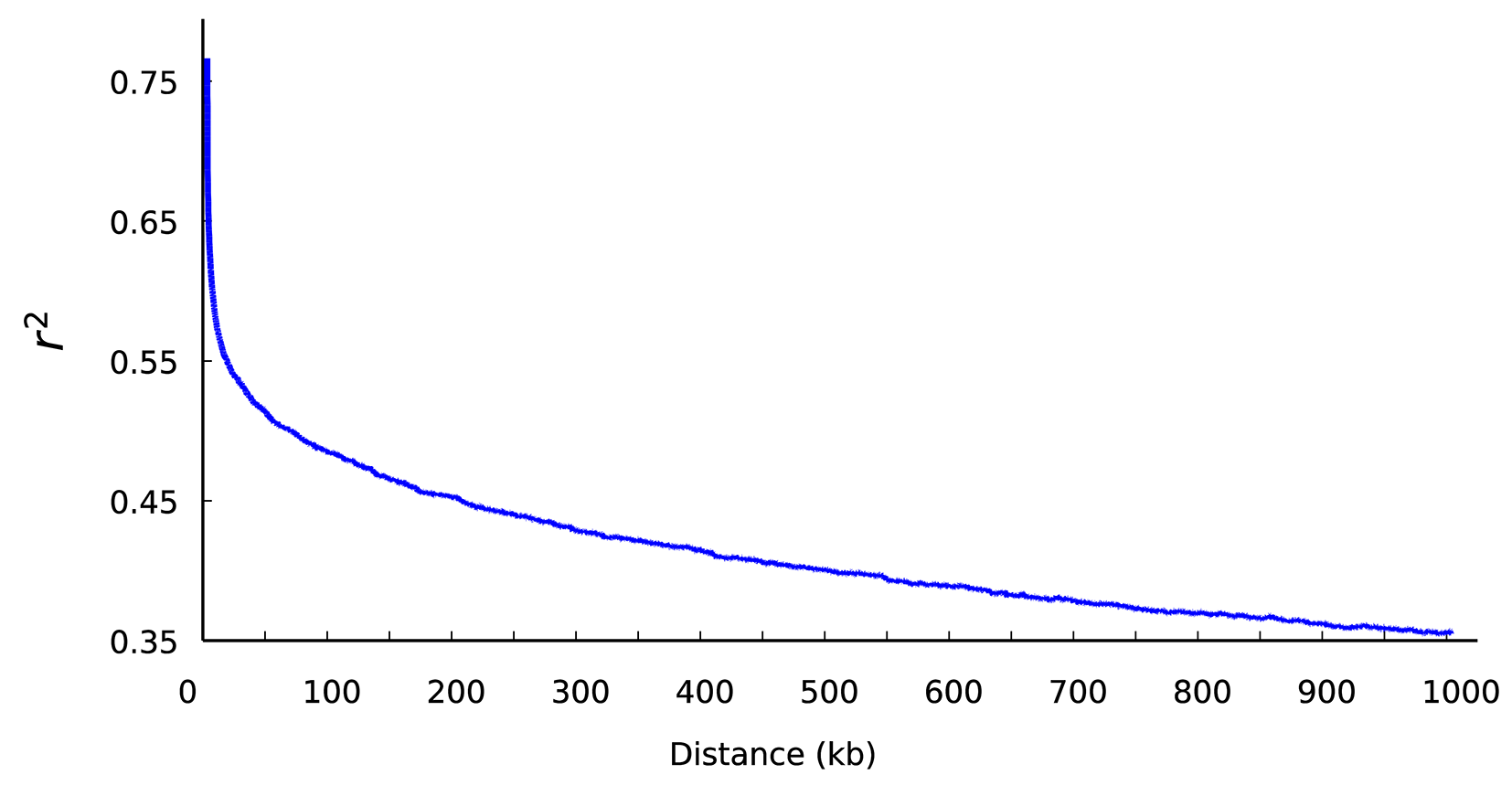

**Supplementary Figure 1**. Decay of linkage disequilibrium (LD) in the snap bean genome

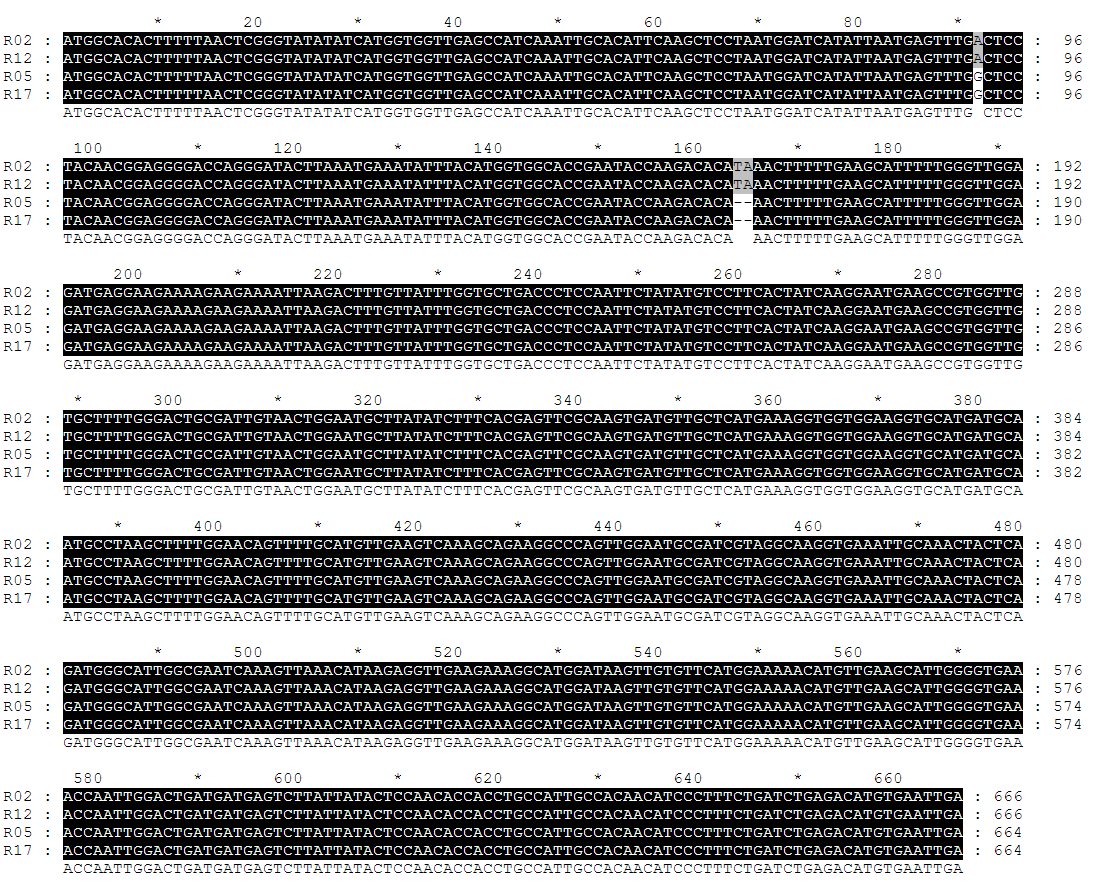

**Supplementary Figure 2**. Sequence alignment of Phvul.002G270800 in four different accessions. R02 and R12 are stringless; R05 and R17 have suture strings.

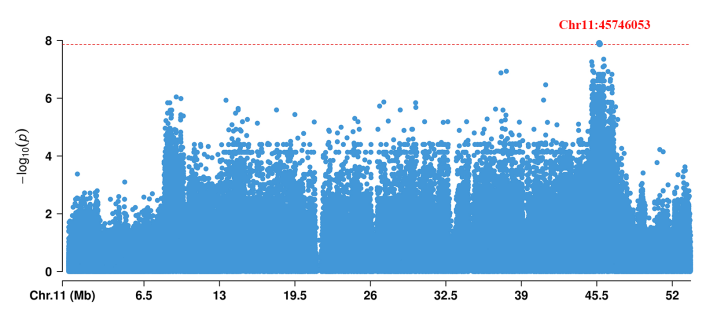

**Supplementary Figure 3.** The Manhattan plots of ST1-2019 on chromosome 11. The red dashed line represents the threshold.

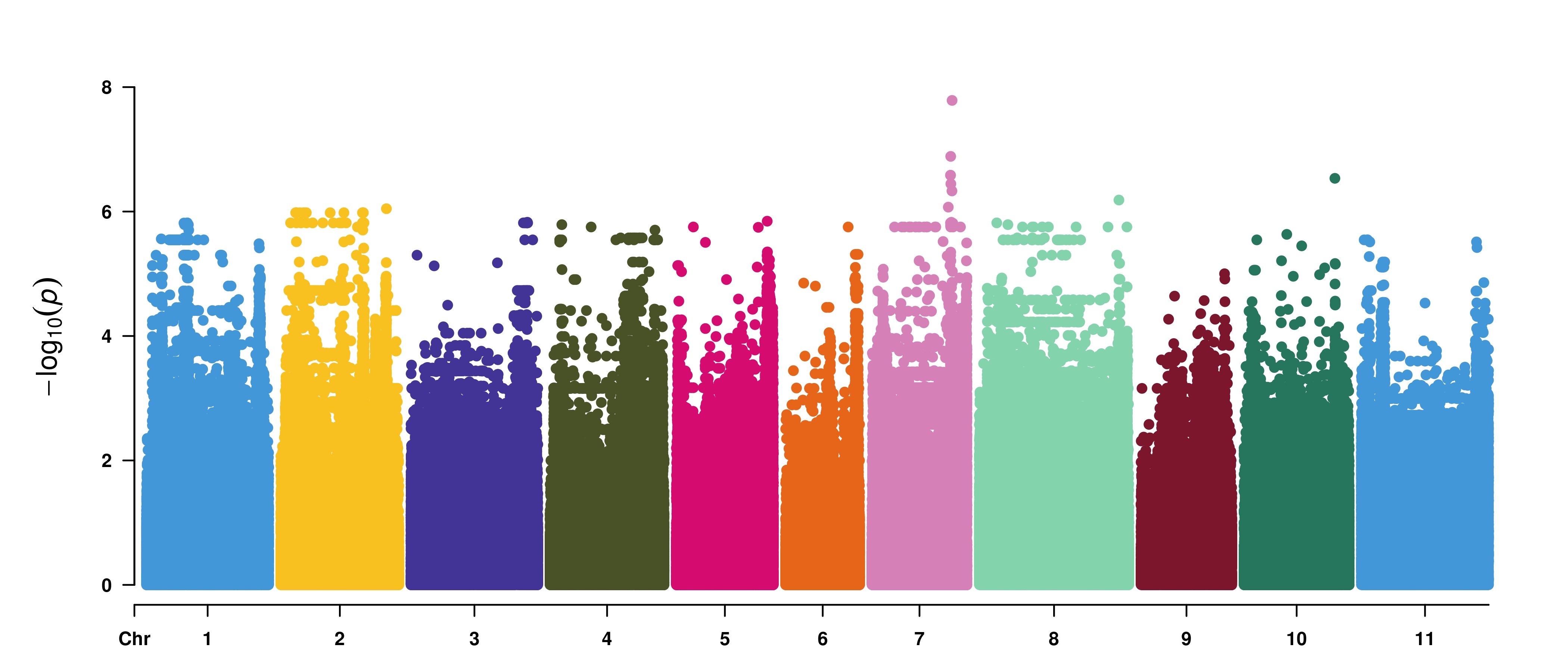

**Supplementary Figure 4.** The Manhattan plots of GWAS for stringy snap bean on ST1-2019
